# Autonomous treatment of bacterial infections *in vivo* using antimicrobial micro- and nanomachines

**DOI:** 10.1101/2021.11.11.468250

**Authors:** Xavier Arqué, Marcelo D. T. Torres, Tania Patiño, Andreia Boaro, Samuel Sánchez, Cesar de la Fuente-Nunez

**Affiliations:** Institute for Bioengineering of Catalonia (IBEC), The Barcelona Institute of Science and Technology (BIST), Barcelona, Spain; Machine Biology Group, Departments of Psychiatry and Microbiology, Institute for Biomedical Informatics, Institute for Translational Medicine and Therapeutics, Perelman School of Medicine, University of Pennsylvania, Philadelphia, Pennsylvania, United States of America; Departments of Bioengineering and Chemical and Biomolecular Engineering, School of Engineering and Applied Science, University of Pennsylvania, Philadelphia, Pennsylvania, United States of America; Penn Institute for Computational Science, University of Pennsylvania, Philadelphia, Pennsylvania, United States of America; Chemistry Department, University of Rome, Tor Vergata, Rome, Italy; Institució Catalana de Recerca i Estudis Avançats (ICREA), Barcelona, Spain

**Author notes:** These authors contributed equally (Xavier Arqué, Marcelo D. T. Torres).

## Abstract

The increasing resistance of bacteria to existing antibiotics constitutes a major public health threat globally. Most current antibiotic treatments are hindered by poor delivery to the infection site, leading to undesired off-target effects and drug resistance development and spread. Here, we describe micro- and nanomachines that effectively and autonomously deliver antibiotic payloads to the target area. The active motion and antimicrobial activity of the silica-based robots are driven by catalysis of the enzyme urease and antimicrobial peptides, respectively. These antimicrobial machines show micromolar bactericidal activity *in vitro* against different Gram-positive and Gram-negative pathogenic bacterial strains and act by rapidly depolarizing their membrane. Finally, they demonstrated autonomous anti-infective efficacy *in vivo* in a clinically relevant abscess infection mouse model. In summary, our machines combine navigation, catalytic conversion, and bactericidal capacity to deliver antimicrobial payloads to specific infection sites. This technology represents a much-needed tool to direct therapeutics to their target to help combat drug-resistant infections.

## INTRODUCTION

Bacterial infections are predicted to kill 10 million people by 2050, which corresponds to one death every three seconds. Today, they are the fourth leading cause of death in hospitals in the USA and cause millions of deaths worldwide, constituting a major public health threat.^1^ Indeed, bacteria are becoming increasingly resistant to broad-spectrum therapies, such as antibiotics, and approaches for countering recalcitrant infections are urgently needed.^2–6^ Currently available antibiotics are limited by their broad and deleterious off-target effects due to the lack of delivery methods that effectively release payloads at the infection site. Advances in nanomedicine have contributed to the development of antibiotic-loaded nanoparticles^7^, but these delivery systems only counter bacterial biofilms at the material interface and can become inert over time due to protein adhesion.^8^ Despite these drastic interventions, infections often remain untreated and untreatable.^9,10^ Hence, novel approaches for effectively delivering antimicrobial payloads to the infection site are urgently needed.^11^

Micro- and nanoparticles can be designed to achieve self-propulsion by converting diverse energy sources into mechanical motion, yielding swimming micro- and nanomachines. Depending on their composition, particles with active motion can be designed for a variety of applications ranging from environmental applications^12^ to biomedicine.^13,14^ Biohybrid active motion has been extensively explored in the past decade in the form of micro- and nano-sized bioactive machines for minimally invasive interventions, given their potential for active navigation to reach otherwise inaccessible areas.^15–17^ Recent breakthroughs have been reported utilizing these systems for biomedical applications, including targeted and enhanced drug delivery,^18,19^ cell manipulation,^20,21^ microsurgery,^22,23^ biochemical sensing,^16,24^ and diagnostics.^25,26^ Bioactive micro- and nanomachines have also been applied to exploit their energy conversion to combat planktonic bacteria^21,27–29^ and biofilms.^30–33^ Indeed, the enhanced mixing and towing force of such moving colloids improves the efficiency of their interaction with bacteria, enabling more effective infection eradication than when using passive particles.^12^ Their navigation increases the probability of contact with the surrounding bacteria and also enhances their penetration capacity.^34^ However, the antibacterial efficacy of micro- and nanomachines has not yet been extensively reported in clinically relevant mouse models,^35–37^ an essential step to eventually translate these applications into the clinic.

In this work, we present bioactive micro- and nanomachines that deliver antimicrobial peptide (AMP) payloads to the infection site through the combination of active motion, catalytic conversion, and bactericidal capabilities. AMPs have emerged as promising antimicrobials^38,39^ whose amphipathic character enables their interaction with and subsequent disruption of bacterial membranes.^40^ Clinical translation of antimicrobials such as AMPs is currently restricted by their limited bioavailability and susceptibility to enzymatic degradation and by their low penetrability towards the target infection.^41^ Thus, efficient delivery methods are required for these molecules to more readily reach their target area.^34^ Delivery vehicles such as unbound liposomes^42^ and nanoparticles^43^ have been used to encapsulate peptides for the treatment of microbial biofilms^42,43^ and cancer,^44^ however these methods rely on passive interactions with the infected area and passive movement through the infection site, *e.g.*, an infected wound.

Here, we used the enzyme urease (a biocompatible source of active motion)^45,46^ to deliver linear, and thus susceptible to proteolytic degradation, cationic AMPs (LL-37 and K7-Pol) on silica-based micro- and nanoparticles that actively navigate in liquid toward the infection site (Figure 1). LL-37 is an amphipathic α-helical and cationic peptide with a 37-amino acid long sequence (LLGDFFRKSKEKIGKEFKRIVQRIKDFLRNLVPRTES). This natural peptide is widely known for its antimicrobial, wound healing, and immunomodulatory properties,^47^ making it a promising candidate for testing in animal models. K7-Pol (ILGTILKLLSKL-NH_2_) is a potent synthetic antimicrobial agent derived from the wasp venom polybia-CP, a mastoparan-like peptide with antimicrobial and immunomodulatory activities ^48^ that presents an amphipathic α-helical and cationic structure and amidated C-terminus. K7-Pol was shown to present broad-spectrum activity against Gram-positive and Gram-negative bacteria and fungi^49^ at low nanomolar concentrations, and against parasites^50^ and cancer cells^51^ at low micromolar concentrations. Despite the potent antimicrobial activity of these AMPs, their stand-alone broad-spectrum activity and susceptibility to protease-mediated degradation leads to off-target effects and reduced activity *in vivo*. Our autonomous machines effectively delivered peptide antibiotic payloads to the infected area, displayed bactericidal activity *in vitro* at low doses (7.8-15.6 μg mL^−1^), targeted bacteria by rapidly depolarizing their membrane, and demonstrated *in vivo* anti-infective efficacy in a clinically relevant mouse model.

**Figure 1.**
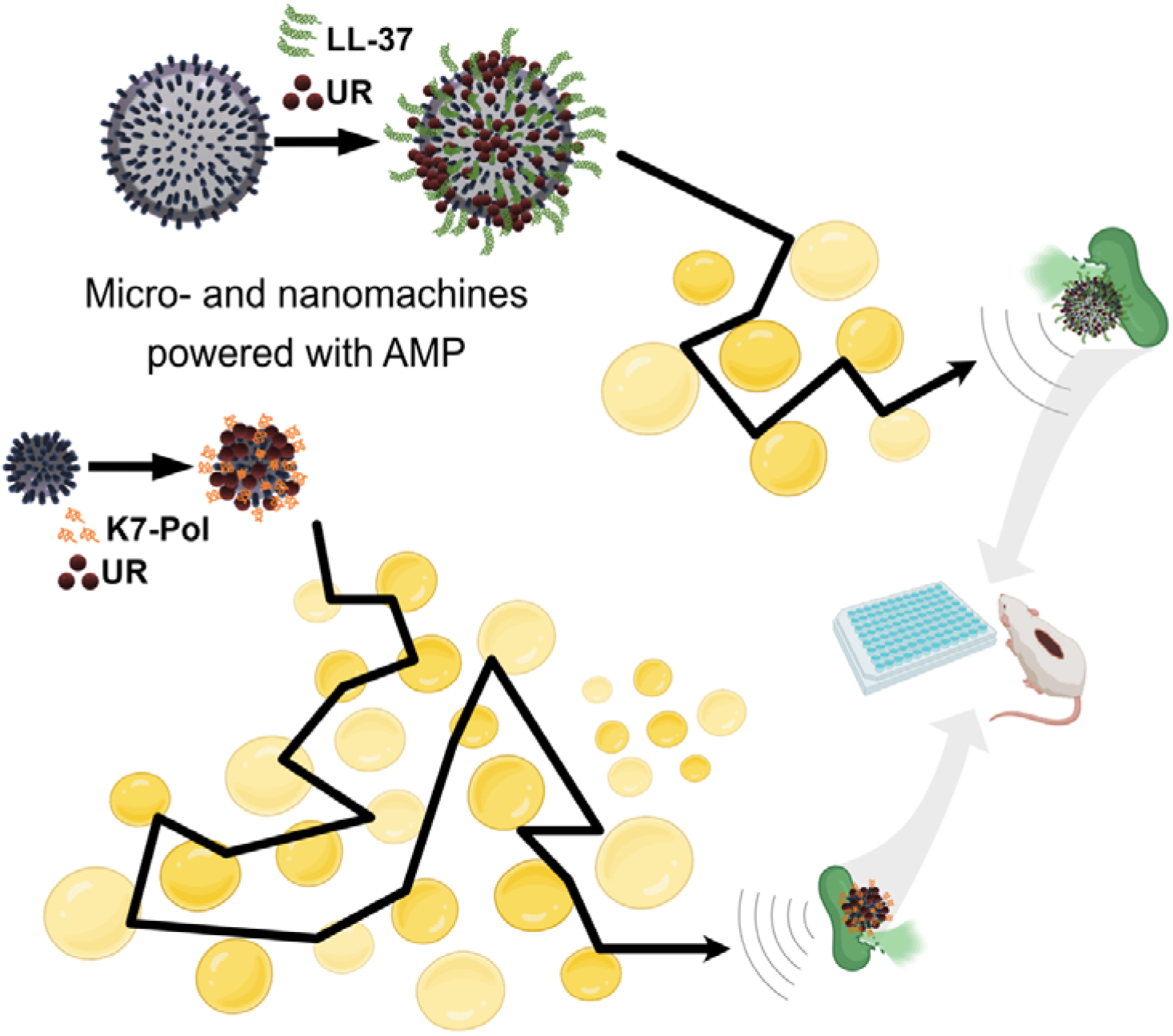
Bioactive micro- and nanomachines coated with antimicrobial peptides for the autonomous treatment of infections. Schematic of the AMP-coating process of the urease micro- and nanomachines and their autonomous propulsion to target pathogenic infections both *in vitro* and *in vivo*. Briefly, AMP-urease machines will encounter and hydrolyze urea in solution. The hydrolysis reaction will propel the machines, and when exposed to bacterial membranes, the AMPs onto their surface will act as antimicrobials and lyse bacterial cells in controlled (*e.g.*, *in vitro* assays) and complex (*e.g.*, infected wound) biological environments.

## RESULTS

### Synthesis and characterization of bioactive micromachines

Bioactive micromachines were fabricated as previously reported^52,53^ through a modified Stöber method^54^ (Figure S1) (see Methods section for details). Briefly, silicon dioxide (SiO_2_) was grown onto commercial 2 μm spherical beads based on polystyrene (PS), using tetraethylorthosilicate (TEOS) and 3-aminopropyltriethoxysilane (APTES) as silica precursors. Later, the PS core was dissolved using N,N-dimethylformamide (DMF) to obtain hollow silica microparticles (HSMP). We characterized the shape, size, and morphology of the microparticles by scanning electron microscopy (SEM), (Figure 2a) and transmission electron microscopy (TEM) (inset Figure 2a) revealing an average microparticle diameter of 1.87 ± 0.01 μm (n = 91, mean ± standard error of the mean). We used TEM to measure the silica particle shell as 48 ± 1 nm (n = 120, mean ± standard error of the mean).

**Figure 2.**
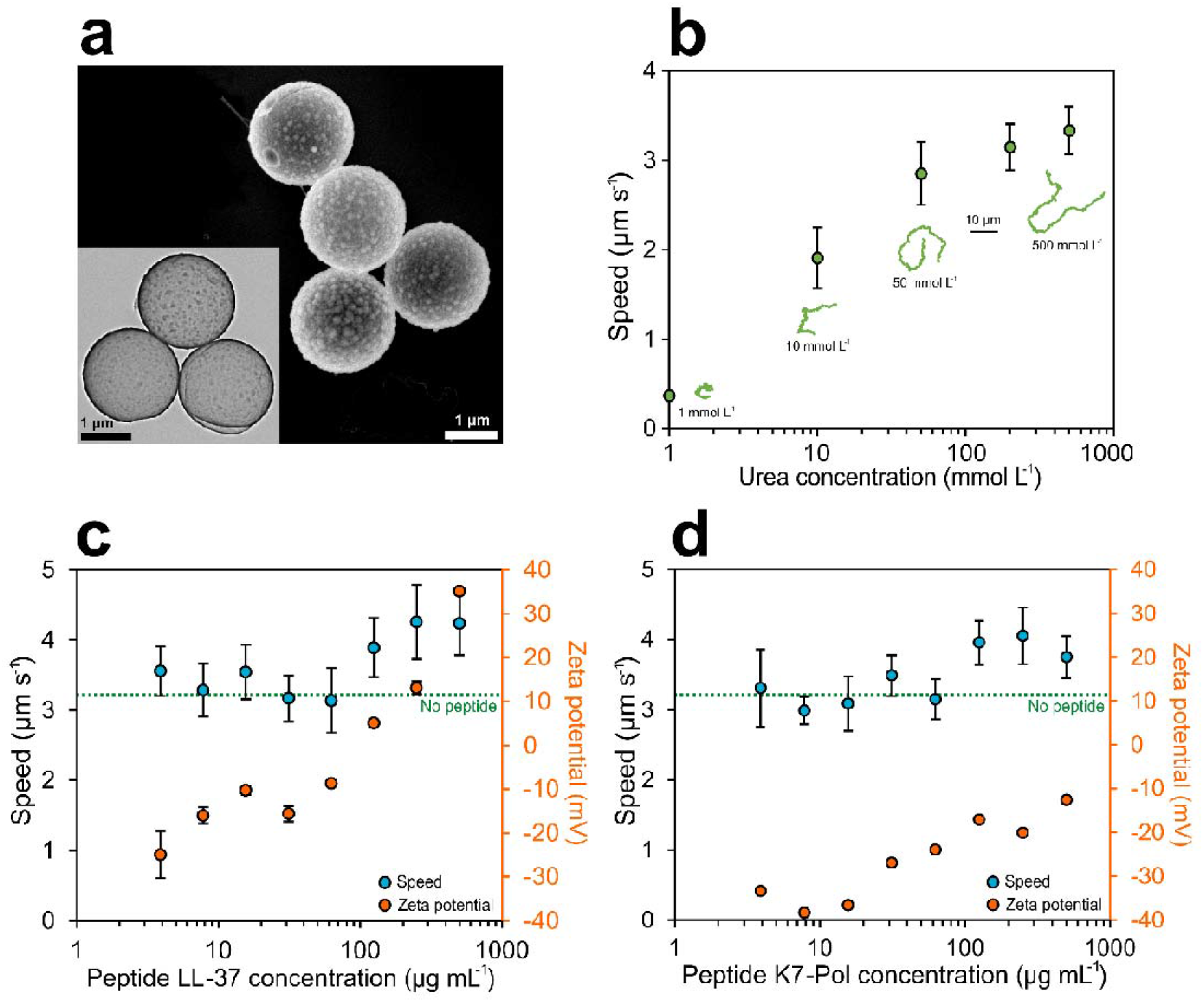
Effect of antimicrobial peptides on the active motion of bioactive micromachines. **(a)** SEM micrograph of the hollow silica microcapsules. Inset: TEM micrograph of the hollow silica microparticles. **(b)** Average speed of urease micromachines for different concentrations of urea. Inset: Representative 15-second trajectories for different concentrations of urea. **(c)** Average speed and zeta potential of urease micromachines for different concentrations of LL-37 peptide used to functionalize the silica surface. **(d)** Average speed and zeta potential of urease micromachines for different concentrations of K7-Pol peptide used to functionalize the silica surface. All results are shown as the mean ± standard error of the mean.

To obtain bioactive micromachines, urease was attached to silica with a glutaraldehyde (GA) linker (Figure 1). For motion experiments, urease micromachines were recorded at 25 FPS for 20-25 seconds. The videos were analyzed using a custom-designed Python-based software in order to extract the trajectories of the micromachines, the mean squared displacement (MSD), and the speed (see Methods section for details).^55,56^ The active motion of urease microparticles was studied as a function of urea (substrate of urease) in water. When the urea concentration increased, the micromachines showed higher MSD (Figure S2) and speed (Figure 2b), which increased in a similar fashion to the Michaelis-Menten saturation curve, as observed in previous publications of urease-powered motion.^53,57^ The self-propulsion capacity reached a plateau with a maximum speed of 3.3 ± 0.3 μm s^−1^ when adding 500 mmol L^−1^ urea. The increment in self-propulsion was clearly distinguished by tracking the trajectories of the individual urease micromachines (Figure 2c, inset and Video 1).

### Preparation of AMP-coated bioactive micromachines

Once the motion behavior of urease micromachines was thoroughly studied and characterized, we incorporated human cathelicidin LL-37 and peptide K7-Pol onto the surface of the silica microparticles (Figure 1). In the last functionalization step to yield the peptide-modified bioactive micromachines, we added both urease and AMP to be anchored to the silica surface by the GA linker (Figure S1) (see Methods section for details). Different concentrations of each peptide were attached to the urease micromachine. By analyzing their electrophoretic mobility through dynamic light scattering (DLS), we observed a considerable increase in the zeta potential (represented by the electrical charge surrounding the particle) when more AMP was added (Figure 2c and d), indicating that the positively charged AMPs were properly coated onto the micro- and nanomachines. For concentrations of LL-37 higher than 125 μg mL^−1^, the microparticle zeta potential changed to positive, reaching a maximum of 35.1 ± 1.4 mV when adding 500 μg mL^−1^ of LL-37 (Figure 2d). In the case of K7-Pol, the zeta potential also increased with higher peptide loads, up to −12.6 ± 0.2 when adding 500 μg mL^−1^ of K7-Pol, but never reached a net positive charge (Figure 2c). The presence of multiple positively charged amino acids in LL-37’s amphipathic sequence (pI = 11.13) (i.e., 5 lysine (K) and 5 arginine (R) residues) explains the drastic change towards a positive zeta potential. In the case of the K7-Pol sequence (pI = 10.6), the presence of 2 lysine (K) residues and the amidated C-terminus also caused an increase in zeta potential but are insufficient to reach a net positive value.

After ensuring the attachment of peptides onto the urease micromachines, we tested their motion using 200 mmol L^−1^ of urea in water as this constituted the lowest urea concentration needed for maximum speed (Fig. 2c). For LL-37-modified micromachines, no significant differences were detected in the urease micromachines’ speed (3.1 ± 0.3 μm s^−1^) when the AMP concentration used was below 62.5 μg mL^−1^ (Figure 2d). Nonetheless, higher doses of AMP (250 μg mL^−1^) caused a significant boost in self-propulsion, increasing speed by 38.7% and reaching 4.3 ± 0.5 μm s^−1^. This increase in directional motion was directly observable by looking at the tracking trajectories extracted from the videos (see Video 2). In the case of K7-Pol-modified micromachines, self-propulsion was also observed when adding K7-Pol, which increased speed by 32.2% for higher concentrations of peptide as indicated by the trajectory tracking experiments (Video 3). In this case, the maximum speed reached was of 4.1 ± 0.4 μm s^−1^ at 250 μg mL^−1^ of K7-Pol (Figure 2d).

While the underlying motion mechanism powered by urease has still not been resolved in the literature, it has recently been pointed out that electric field generation by the release of ionic products could play a significant role.^52,58^ In this regard, the fact that higher concentrations of peptide led to higher propulsion capabilities could be explained by an increase in the surface net charge of the particle, leading to higher conductivity and enhanced electric field generation.

### Synthesis and characterization of bioactive nanomachines

Mobil Composition of Matter No. 41 (MCM-41) mesoporous silica nanoparticles (MSNP) were chosen as a base material for the bioactive nanomachines because of their biocompatibility and ease of surface modification,^59,60^ as well as their extensive use for biomedical applications.^61^ Urease nanomachines were fabricated as recently reported^34,62^ using a modified Stöber method^63^ (Figure S3) (see Methods section for details). The nanoparticles were prepared by hydrolysis and condensation of TEOS in aqueous media basified with triethanolamine (TEOA) and using hexadecyltrimethylammonium bromide (CTAB) as the structure-directing agent. Next, to obtain the mesoporous silica nanoparticles, the CTAB surfactant was removed by acidifying the solution in methanol. The generated MSNP were then treated with APTES to modify the silica surface with amine groups. The morphology of the nanoparticles was characterized by both SEM and TEM (Figure 3a), displaying an average diameter of 694.698 ± 0.003 nm (n = 63, average size ± standard error of the mean). A clear radial mesoporosity was observed when increasing the magnification of TEM micrographs (Figure S4). The amino groups on the surface of the MSNP were later used to attach the urease enzyme through the GA linker to yield urease nanomachines.

**Figure 3.**
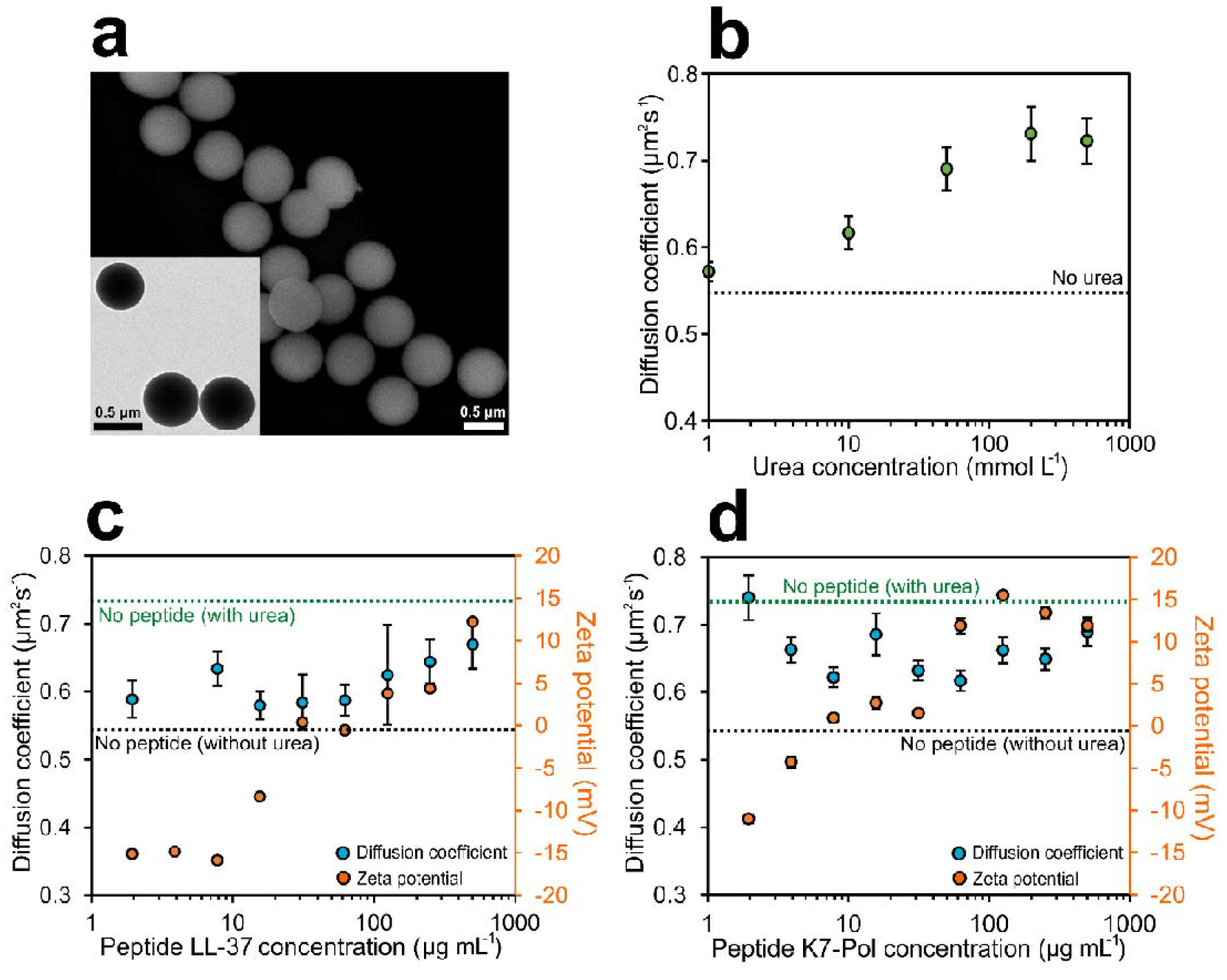
Effect of antimicrobial peptides on the active motion of bioactive nanomachines. **(a)** SEM micrograph of the mesoporous silica nanoparticles. Inset: TEM micrograph of the mesoporous silica nanoparticles. **(b)** Diffusion coefficient of urease nanomachines for different concentrations of urea. **(c)** Diffusion coefficient and zeta potential of urease nanomachines for different concentrations of LL-37 peptide used to functionalize the silica surface. **(d)** Diffusion coefficient and zeta potential of urease nanomachines for different concentrations of K7-Pol peptide used to functionalize the silica surface. All results are shown as the mean ± standard error of the mean.

For the motion experiments, we studied the electrophoretic mobility of urease nanomachines through DLS to obtain both their diffusion coefficient and apparent hydrodynamic radii (see Methods section for details). The active motion of urease nanomachines was studied as a function of urea concentration in PBS, and for all conditions a sharp single population peak was observed (Figure S5), confirming monodispersity of the sample. The diffusion of nanomachines was enhanced when increasing the urea concentration (Figure 3b), from a base Brownian diffusion of 0.54 ± 0.01 μm^2^ s^−1^ when no substrate was present, to a maximum enhanced diffusion of 0.73 ± 0.03 μm^2^ s^−1^ at 200 mmol L^−1^ urea. This 35.2% increase in diffusion as a function of urea concentration, and the further saturation of active motion resemble the dynamics of a Michaelis-Menten kinetics curve, as previously reported for urease-powered enhanced diffusion.^18,34,62,64^ The apparent hydrodynamic radii decreased from 451.7 nm to a minimum of 329.2 nm at 200 mmol L^−1^ urea, since when increasing the diffusion coefficient (D) with more urea the radii values obtained though the Stokes−Einstein equation also decreased accordingly to match the diffusion measured (Figure S5) (see Methods section for details).

### Preparation of AMP-coated bioactive nanomachines

Upon determining the motion behavior of urease nanomachines, the fabrication process was modified to incorporate the human cathelicidin peptide LL-37 and the potent synthetic antimicrobial peptide K7-Pol onto the surface of the silica nanoparticles (Figure 1).

For the last functionalization step, we added both urease and AMP to the solution containing nanoparticles, to enable binding of urease and the AMP to the GA linker onto the silica surface (see Methods section for details). A range of AMP concentrations were attached to the urease nanomachines, and analysis of their electrophoretic mobility using DLS revealed a considerable increase in the zeta potential (represented by the electrical charge surrounding the particle) with increasing levels of peptide (Figure 3c and d). A positive zeta potential of the nanomachines was reached by adding 31.25 μg mL^−1^ of LL-37 and a maximum of 12.3 ± 0.2 mV was measured at 500 μg mL^−1^ of LL-37 (Figure 3c). In the case of K7-Pol, at least 7.81 μg mL^−1^ of K7-Pol were required to obtain a positive zeta potential (Figure 3d), reaching a maximum of 15.5 ± 0.2 mV at 125 μg mL^−1^ of K7-Pol which was saturated for higher AMP concentrations. These data are consistent with the presence of 2 lysine residues (K) and the amidated C-terminus of the K7-Pol sequence, yielding a pI of 10.6, and the presence of five lysine (K) and five arginine (R) residues in LL-37’s amphipathic sequence with a pI of 11.13.

Once we confirmed the attachment of the peptides to the urease nanomachines, we tested their motion capabilities at 200 mmol L^−1^ urea in PBS, as this was the urea concentration that allowed maximum diffusion. The diffusion of LL-37-coated nanomachines decreased when attaching the peptide, however the particles still showed active motion under all conditions tested (Figure 3c and d). The lowest diffusion was 0.58 ± 0.02 μm^2^ s^−1^ at 15.62 μg mL^−1^ LL-37, representing a 78.9 % reduced diffusion and there were no significant differences in diffusion at the different AMP concentrations tested (Figure 3c). Figure 3d shows that attachment of K7-Pol onto the nanoparticle surface also resulted in decreased enhanced diffusion at concentrations higher than 1.95 μg mL^−1^ (Figure 3d). The maximum decrease in enhanced diffusion was 61.3 %, from 0.73 ± 0.03 μm^2^ s^−1^ without peptide to 0.62 ± 0,02 μm^2^ s^−1^ at 62.5 μg mL^−1^ K7-Pol. Hence, the enhanced diffusion decreased when attaching AMPs is likely due to direct competition of the peptides with urease for the available functional groups on the silica surface, causing a decrease in both enzyme attachment and active motion. However, for the purpose of fabricating active antimicrobial tools, it is important to note that the peptide-urease nanomachines still presented active motion for all the different peptide concentrations tested.

### *In vitro* antimicrobial activity of micro- and nanomachines against pathogenic bacteria

The AMP-modified bioactive micro- and nanomachines were tested against bacteria from the ESKAPE pathogen list declared by the World Health Organization as critical threats for humans: *Acinetobacter baumannii* AB177, *Escherichia coli* ATCC11775, *Klebsiella pneumoniae* ATCC13883, *Pseudomonas aeruginosa* PAO1, and *Staphylococcus aureus* ATCC12600. Minimum inhibitory concentration (MIC) values were determined as the lowest concentration of our machines tested in the presence of 200 mmol L^−1^ urea that inhibited 100% of bacterial growth. The MIC values for each condition were assessed by determining the optical density of the solution at 600 nm and ranged from 7.8 to 125 μg mL^□1^ depending on the bacterial strain (Figure 3). Both micro- and nanomachines exhibited 4- to 16-fold enhanced antimicrobial activity when the AMPs were incorporated (Figure 3). Interestingly, AMP-modified machines displayed activity against *K. pneumoniae* and *S. aureus*, whereas neither the bare micro-nor the nanomachines presented activity toward *S. aureus*, and the bare nanomachines were inactive against *K. pneumoniae*.

### Mechanism of action studies

Mechanism of action (MoA) studies were carried out to explore whether AMP-modified micro- and nanomachines killed bacterial cells by permeabilizing their outer membrane or, instead, acted by depolarizing their cytoplasmic membrane at their MIC. To assess if the AMP-modified bioactive machines were able to permeabilize the outer membrane of *A. baumannii* and *K. pneumoniae* cells, we used the fluorescent probe NPN [1-(N-phenylamino)-naphthalene]. NPN exhibits weak fluorescence emission in aqueous environments and can permeate the bacterial outer membrane when damaged. The probe interacts with the lipidic environment of permeable outer membranes and displays an increased intensity of fluorescence emission (Figure S6). As a positive control, we used the FDA-approved and widely used peptide antibiotic polymyxin B (PMB), a well-known membrane permeabilizer. When we exposed *A. baumannii* or *K. pneumoniae* cells to the AMP-modified micro- and nanomachines and PMB, only PMB was able to permeabilize the outer membrane (Figure S6).

To evaluate whether the AMP-urease micro- and nanomachines were able to depolarize the cytoplasmic membrane of *A. baumannii* and *K. pneumoniae* cells, we used the probe DiSC_3_-5 (3,3’-dipropylthiadicarbocyanine iodide). DiSC_3_-5 is a potentiometric probe that accumulates in the cytoplasmic membrane and aggregates at high concentrations causing fluorescence quenching. ^65,66^ When the cytoplasmic membrane is depolarized, DiSC_3_-5 migrates to the cytoplasm, leading to increased fluorescence emission intensity (Figure 5a). The micro- and nanomachines with and without AMP had similar ability to depolarize the cytoplasmic membrane of *A. baumannii* (Figure 5b). However, the AMP-urease machines presented increased depolarization of the *K. pneumoniae* membrane than their bare counterparts (Figure 5c). This result is in accordance with the observed depolarizing effect of the peptides.^67^ K7-Pol-urease nanomachines showed a slightly increased depolarization effect than LL-37-urease nanomachines. This was not entirely unexpected, since mastoparans, such as K7-Pol and all polybia-CP-derived peptides, depolarize bacterial membranes more efficiently than defensins (*e.g.*, LL-37).^65^ All experiments were performed in the presence of 200 mmol L^−1^ urea.

### Anti-infective activity against *A. baumannii* in a skin abscess animal model

All the combinations of micro- and nanomachines and peptides, and their bare urease micro- and nanomachines counterparts were tested *in vivo* against *A. baumannii* at 2-fold their MIC, 15.6 and 31.2 μg mL^□1^, respectively (Figure 6a). Skin infection was induced by administering an *A. baumannii* solution at 10^7^ CFU mL^□1^ on the back of mice previously scratched with a needle. ^65,68,69^ The treated groups were administered with a single dose of free AMPs, micro- or nanoparticles (with and without AMP) two hours post-infection. The samples were added to one extremity of the infected wound and 100 μL of a 200 mmol L^−1^ urea solution was spread over the entire length of the wound (Figure 6b). Four days after a single treatment dose, the most active systems were LL-37-urease micromachines and K7-Pol-urease nanomachines, significantly reducing the bacterial load by 2 and 3 orders of magnitude, respectively (Figure 6c), to levels that can be cleared up by the immune response (<10^3^ CFU mL^−1^).

On the contrary, treatment with AMPs free in solution (Figure 6b) that were administered in one of the extremities of the wound only displayed anti-infective activity locally where the peptides were administered (Figure 6d). More specifically, peptide treatment reduced CFU counts by 2-3 orders of magnitude in the administration site (~1/3 of the whole extent of the wound), whereas infected areas farther away from the administration site remained infected at levels similar to the untreated controls. Interestingly, for LL-37, which presents immunomodulatory and wound healing properties, ^70^ we also observed partial wound healing in the area where the peptides were added (Figure S7). To assess whether the peptides free in solution and the AMP-modified bioactive micro- and nanomachines were toxic to the animals, the weight of the mice was monitored throughout the experiment. Variations of up to 20% in weight are a widely used proxy of distress, morbidity, and overall toxicity ^49,69^. No side effects (*e.g.*, itchiness, redness, swelling) or *in vivo* toxicity were observed under any of the conditions tested (Figure 6e-f). The LL-37-urease nanomachines and K7-Pol-urease micromachines and the machines without peptide presented bacteriostatic effects (Figure 4).

**Figure 4.**
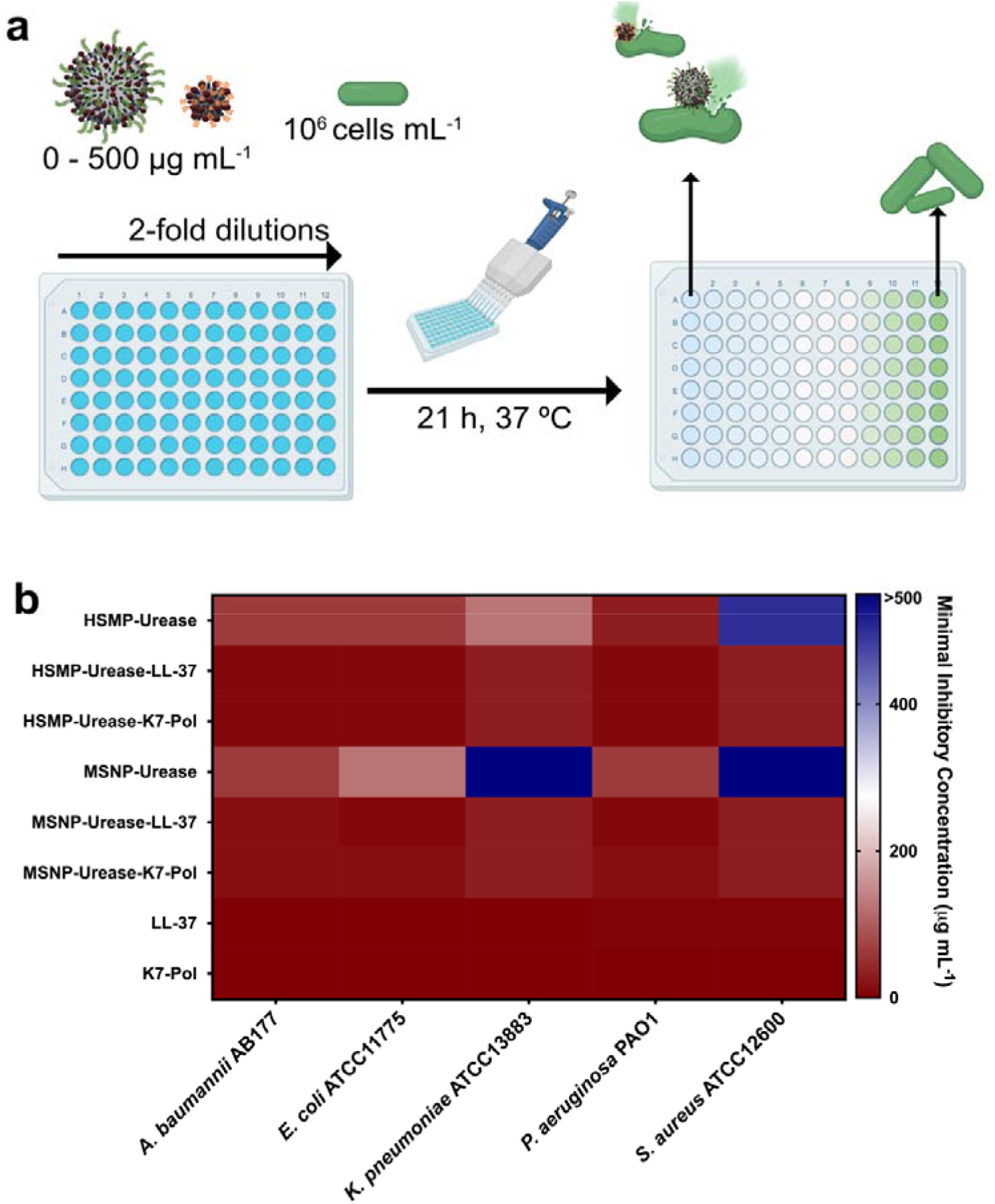
Antimicrobial activity of bioactive micro- and nanomachines functionalized with antimicrobial peptides. **(a)** Schematic of the *in vitro* biological activity experimental design. Briefly, 10^5^ bacterial cells and urease micro- and nanomachines (0-500 μg mL^−1^) were added to a 96-wells plate and incubated at 37 °C. One day after the exposure, the solution in each well was measured in a microplate reader (600 nm) to check inhibition of bacteria compared to the untreated controls. **(b)** Heat map of the antimicrobial activity of each system against five bacterial strains: *A. baumannii* AB177, *E. coli* ATCC11775, *K. pneumoniae* ATCC13883, *P. aeruginosa* PAO1, and *S. aureus* ATCC12600. Assays were performed in three independent replicates and heat map OD_600_ values are the arithmetic mean of the replicates in each condition.

**Figure 5.**
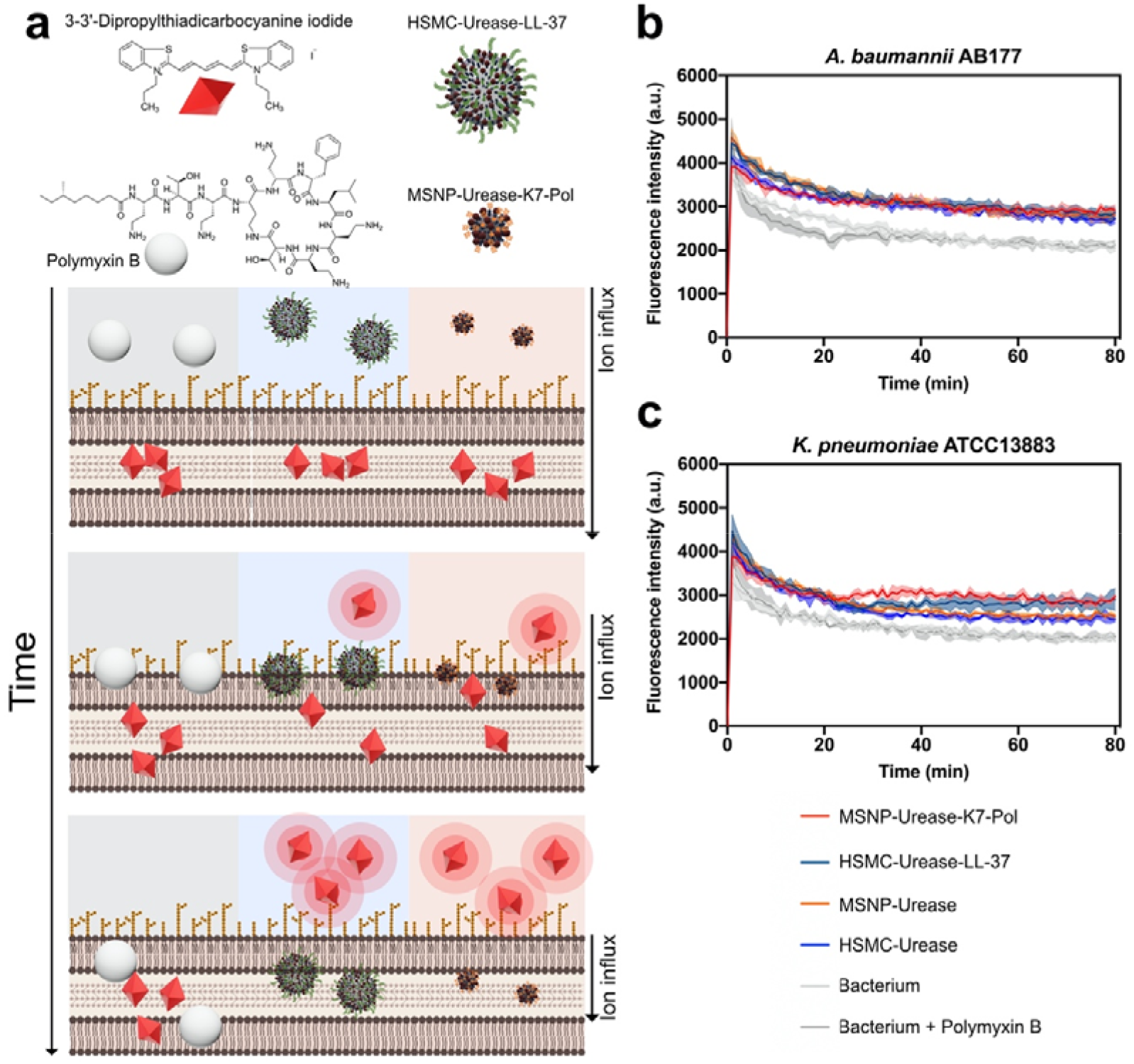
Mechanism of action of the antimicrobial machines. **(a)** Bioactive micro- and nanomachines cause the depolarization of bacterial membranes at their MIC concentration against **(b)** *A. baumannii* AB177 and **(c)** *K. pneumoniae* ATCC13883. Briefly, micro- and nanomachines functionalized with LL-37 and K7-Pol, respectively, enabled the higher depolarization of *K. pneumoniae* cells than the non-functionalized machines. When *A. baumannii* cells were exposed to them both functionalized and non-functionalized machines presented depolarizing effect. The potent permeabilizer antimicrobial polymyxin B was used as a negative control for depolarization.

**Figure 6.**
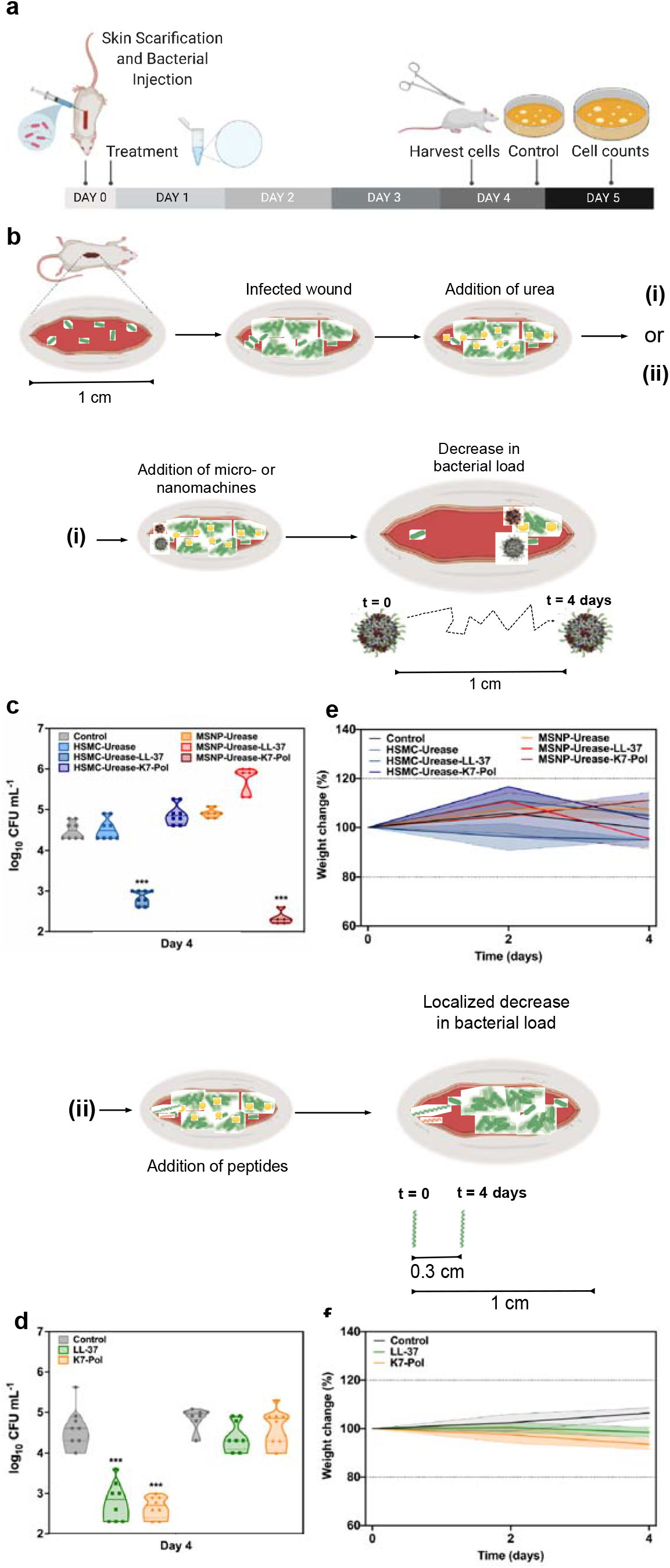
Anti-infective activity of the antimicrobial machines *in vivo*. **(a)** Mice had their dorsal region shaved, scratched (1 cm-long wound) and infected with *A. baumannii*. After one hour, functionalized and non-functionalized micro- and nanomachines or peptides were added to the infection site. Mice were euthanized and the tissue from the infection site was harvested, and the bacterial cells of treated and untreated samples were counted by plating. **(b)** Schematic representation of the wound site infected and the addition of urea before the treatment with (i) micro- and nanomachines or (ii) peptides free is solution. Antimicrobial micro- and nanomachines self-propelled, driven by urea, through a distance of 1 cm to enable the autonomous treatment of the target infected area. On the other hand, peptides by themselves only exhibited antimicrobial activity within the area they were administered and did not clear the infection at a distance. Briefly, after the infection was established, urea was spread over the entire length of the wound (1 cm). Next, the micro- and nanomachines coated with peptides and the peptides alone were inoculated to one of the extremities of the infected wound. **(c)** Four days post-infection, 1 cm^2^ of the infected area was excised and the ability of the micro- and nanomachines to travel throughout the wound alone and when functionalized with peptides was assessed. **(d)** Treatment with peptides alone decreased bacterial counts only in the extremity where they were administered (light yellow background) as revealed by similar bacterial counts detected in areas at a distance from the administration site (dark yellow background) and those of untreated control groups. **(e)** Mouse weight was monitored throughout the experiments, serving as a proxy to assess the toxicity of both micro- and nanomachines and **(f)** peptides in solution. None of the treatment groups led to toxicity in mice.

To shed light into the underlying mechanisms associated with the differences in antimicrobial activity observed between our *in vitro* and *in vivo* experiments, we investigated the speed of each of the machines. No significant differences were observed when comparing the speed of both AMP-modified micromachines, nor when comparing the diffusion coefficient of both AMP-modified urease nanomachine (Figure S8). Thus, we hypothesize that LL-37-urease micromachines and K7-Pol-urease nanomachines presented activity in the animal model because less peptide is needed to obtain more positively charged systems for those combinations. LL-37-urease micromachines presented zeta potential values of −10.3 ±1.9 while K7-Pol-urease micromachines showed −36.6 ±0.1 at the concentration used in the animal model (15.6 μg mL^□1^). On the other hand, K7-Pol-urease nanomachines presented zeta potential values of 1.5 ±0.2, whereas LL-37-urease nanomachines exhibit a zeta potential value of 0.4 ±0.2 at the concentration used *in vivo* (31.2 μg mL^□1^) (Figures 2 and 3). These values indicate that the initial electrostatic interactions between the negatively charged bacterial membranes and the AMP-urease machines are likely crucial to trigger depolarization and subsequent bacterial death.

## CONCLUSION

We report the fabrication and detailed characterization of urease micro- and nanomachines that actively deliver peptides to combat clinically relevant infections. The motion dynamics of each particle was studied under different concentrations of urea and active motion was confirmed upon functionalization with the antimicrobial peptides. The antimicrobial micro- and nanomachines displayed broad-spectrum bactericidal activity *in vitro* against both Gram-negative (*A. baumannii*, *E. coli*, *K. pneumoniae*, *P. aeruginosa*) and Gram-positive (*S. aureus*) bacterial pathogens. An in-depth mechanistic study confirmed that both bare urease micro- and nanomachines targeted bacteria by depolarizing their membrane and coating them with antimicrobial peptides led to enhanced bactericidal activity and bacterial membrane depolarization. Finally, the AMP-modified micro- and nanomachines demonstrated autonomous propulsion to reduce *A. baumannii* infections by up to 3 orders of magnitude in a preclinical murine infection model, whereas free peptides in solution were not able to exert antimicrobial activity at a distance from the initial administration site. We show that differences in surface net positive charge of the micro- and nanomachines coated with peptides influenced antimicrobial activity *in vivo* by increasing the activity range along the wound, but not in less complex environments (*in vitro*). This work constitutes a relevant step toward the implementation of bioactive and autonomous micro- and nanomachines for the treatment of infectious diseases.

## Methods

### Chemicals

2□μm microparticles based on polystyrene (Sigma-Aldrich cat. no. 78452), ethanol 99% (PanReac AppliChem cat. no. 131086-1214), ammonium hydroxide solution 28-30% (Sigma-Aldrich cat. no. 221228), 3-aminopropyltriethoxysilane (APTES) 99% (Sigma-Aldrich cat. no. 440140), triethanolamine (TEOA) 99% (Sigma-Aldrich cat. no. 90279), hexadecyltrimethylammonium bromide (CTAB) 99% (Sigma-Aldrich cat. no. 52365), methanol (MeOH) 99% (PanReac AppliChem cat. no. 361091-1611), hydrochloric acid (HCl) 37% (PanReac AppliChem cat. no. 211020-1611), tetraethylorthosilicate (TEOS) 99% (Sigma-Aldrich cat. no. 86578), N,N-dimethylformamide (DMF) 99% (Acros Organics cat. no. 423640010), 1x phosphate-buffered saline (PBS) (Thermo Fisher Scientific cat. no. 70011-036), glutaraldehyde (GA) (25□wt%) (Sigma-Aldrich cat. no. G6257), urease from *Canavalia ensiformis* (Jack bean) (Sigma-Aldrich cat. no. U4002), urea (Sigma-Aldrich cat. no. U5128).

### Instruments

SEM images were captured by a FEI NOVA NanoSEM 230. TEM images were captured by a Zeiss EM 912. The zeta potential (ζ potential) measurements were performed with a Zetasizer Nano S from Malvern Panalytical. The hydrodynamic radius and diffusion coefficient measurements were performed using a Möbius from Wyatt Technology. The optical videos of urease micromachines were recorded using the camera (Hamamatsu Digital Camera C11440) of an inverted optical microscope (Leica DMi8). The optical density (OD) of the antibacterial assays was measured at 600 nm in a Thermo Scientific Varioskan LUX fluorescence spectrophotometer.

### Antimicrobial peptide synthesis

The peptides LL-37 and K7-Pol were purchased from Aapptec (Kentucky, USA). They were purified by high-performance liquid chromatography (HPLC). The peptide purity used in all assays was higher than 95%.

### Synthesis of hollow silica microparticles (HSMP)

The HSMPs were synthesized by mixing 500□μL of 2□μm spherical beads based on polystyrene (PS) (Sigma-Aldrich cat. no. 78452), 1□mL ethanol 99% (Panreac Applichem cat. no. 131086-1214), and 0.8[ml ultrapure water. Next, 50□μL ammonium hydroxide 28-30% (Sigma-Aldrich cat. no. 221228) was added and the solution was let to magnetically stir for 5□min. Then, 5□μL 3-aminopropyltriethoxysilane (APTES) 99% (Sigma-Aldrich cat. no. 440140) was added, and the reaction was mixed for the following 6 h. After, 15□μL tetraethylorthosilicate (TEOS) ≥99% (Sigma-Aldrich cat. no. 86578) was added to the solution, and the reaction was let to continue stirred overnight (17 h). Next, the PS beads coated with silica were washed 3 times with ethanol (centrifugation of microparticles is always at 1503□rcf for 4 min). The PS was then dissolved from inside the silica shell with 4 washes of dimethylformamide (DMF) ≥99.8% (Acros Organics cat. no. 423640010), with a 15-min mixing step in each wash. Afterwards, the HSMPs obtained were washed 3 times with ethanol 99% and stored at room temperature.

### Functionalization of HSMPs with urease and AMPs

To fabricate silica micromachines, the HSMP were washed 3 times with ultrapure water and 1 time with 1x phosphate-buffered saline (PBS) (pH□=□7.4) (Thermo Fischer Scientific cat. no. 70011-036). Then, the particles were suspended in 1x PBS containing glutaraldehyde (GA) at 2.5□wt% (Sigma-Aldrich cat. no. G6257) and kept mixing at room temperature for 3□h. Next, the silica microparticles functionalized with GA were washed 3 times with 1x PBS (pH□=□7.4) and resuspended again in 1x PBS (pH□=□7.4) with 3□mg□mL^−1^ of urease powder from *Canavalia ensiformis* (Jack bean) (Sigma-Aldrich cat. no. U4002). When fabricating micromachines with peptides attached, different concentrations of AMP (500, 250, 125, 62.5, 31.25, 15.62, 7.81, 3.9 and 0 μg mL^−1^) were added simultaneously with urease. The solution was kept mixing overnight (16 h) and then washed 3 times with 1x PBS (pH□=□7.4). Then, the solution of urease micromachines in 1x PBS (pH□=□7.4) was divided in aliquots and stored at 4 °C to be used for further experiments.

### Optical video recording of bioactive micromachines motion

To study the active motion of urease micromachines, optical videos were recorded using a digital camera (Hamamatsu Digital Camera C11440) coupled with an inverted optical microscope (Leica DMi8). The 63x water immersion objective was used to record the micromachines placed on a glass slide, thoroughly mixed with the water solutions of urea (enzymatic substrate) at different concentrations (500, 200, 50, 10, 1 and 0 mmol L^−1^), specifically selected to cover the range at which urease was active and showed the Michaelis-Menten kinetics saturation, as reported in BRENDA, the Comprehensive Enzyme Information System (https://www.brenda-enzymes.org/). For the motion experiments in function of different AMP concentration (500, 250, 125, 62.5, 31.25, 15.62, 7.81, 3.9 and 0 μg mL^−1^), 200 mM urea to ensure optimal catalytic rate and active motion. The micromachines solution was put on a glass slide covered with a coverslip and videos of 25□FPS and 20-25□s were recorded up to the first 3□min after mixing. For each condition studied of urea and AMP concentration, 10-14 individual urease micromachines were recorded.

### Data analysis of bioactive micromachine motion

The videos were analyzed using custom-designed tracking Python software to obtain the tracking trajectories of the microparticle displacement. From the *X* and *Y* axes values over time, the MSD was calculated using the following equation:

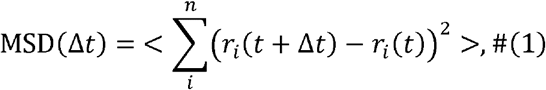

where *t* is the time, *r*_*i*_(*t*) is the position of the particle in the coordinate *i* at time *t*, *n* = 2 are the dimensions 2D analysis, and <·> denotes ensemble and time average. The velocity (*v*) was then extracted from fitting the MSD to

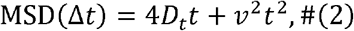

where *D*_*t*_ is the diffusion coefficient and *v* is the speed, since we analyze the propulsive regime when *t* ≪ *τ*_*r*_, being *τ*_*r*_ the rotational diffusion time, and *t* the time of MSD represented.^55,56^ The results are presented as mean ± standard error of the mean.

The *τ*_*r*_ was calculated to be 4.575 ± 0.044 s, which is

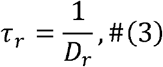

where *D*_*r*_ or is the rotational diffusion coefficient (*D*_*r*_ = 0.219 ± 0.002 s^−1^), that depends on the radius of the particle, as it can be observed in the Stokes-Einstein equation,

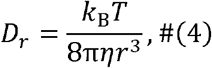

where *k*_*B*_ is the Boltzmann constant, *T* is the absolute temperature, *η* is the solvent viscosity and *r* is the radius of the diffusing particle. Hence, *τ*_*r*_ depends on the temperature (*T* = 24 ± 1 °C), the solvent viscosity (*η* = 0.9107·10^−3^ kg m^−1^ s^−1^), and the radius of the particle (*r* = 0.936 ± 0.003 μm). A nano/micromotor analysis tool was used to compile and visualize the resulting files (https://github.com/rafamestre/NMAT-nanomicromotor-analysis-tool).

### Synthesis of Mesoporous Silica Nanoparticles (MSNP)

The MSNPs were synthesized through sol−gel methodology.^63^ Briefly, 570 mg CTAB (Sigma-Aldrich cat. no. 52365) and 35 g TEOA (Sigma-Aldrich cat. no. 90279) were dissolved in 20 mL of ultrapure water and heated at 95 °C in a silicon oil bath. This solution was stirred for 30 min, after which 1.5 mL of TEOS (Sigma-Aldrich cat. no. 86578) was added dropwise. The solution was then stirred at 95 °C for 2 h stirring in a silicon oil bath. After, the resulting particles were collected by centrifugation and washed 3 times with ethanol (centrifugation of nanoparticles is always at 1503 rcf for 5 min). Then, to remove the CTAB from the MSNP pores, the particles were suspended in a solution composed by 30 mL methanol (PanReac AppliChem cat. no. 361091-1611) and 1.8 mL HCl (PanReac AppliChem cat. no. 211020-1611) and left in the reflux at 80 °C for 24 h. After collecting the particles by centrifugation, 3 washes of ethanol were applied with 10 min of sonication for each wash. Finally, 3 aliquots of 0.5 mL were taken from the resulting solution to centrifuge them and determine the concentration by measuring the weight before and after air-drying the solvent.

### Amino modification of the MSNP surface

The silica surface of the MSNPs was modified with APTES to incorporate amino groups, by adjusting an already reported method.^71^ First, a solution of 2 mg mL^−1^ of MSNP was prepared in a round bottom flask and heated up at 50 °C under magnetic stirring. Next, APTES was added to stirring solution to obtain a final concentration of 5 mM, and it was left under reflux at 50 °C for 24 h. After this process, the resulting MSNPs-NH_2_ were washed 3 times with ultrapure water and 3 aliquots of 0.5 mL were taken to determine the concentration by measuring the weight before and after air-drying the solvent inside.

### Functionalization of MSNP-NH2 with urease and AMPs

The resulting MSNP-NH_2_ were diluted in ultrapure water to obtain a 1 ml solution of 1 mg mL^−1^ of nanoparticles, then centrifuged, resuspended in 900 μL of PBS (pH□=□7.4) and sonicated for 10 min. After this, 100 μL of GA at 25□wt% was added and left to mix for 2.5 h at room temperature. The MSNP-NH_2_ were then washed 3 times with PBS, applying a 10-minute sonication for each wash. Next, the silica nanoparticles functionalized with GA were washed 3 times with 1x PBS (pH□=□7.4) and resuspended again in 1x PBS (pH□=□7.4) with 3□mg□mL^−1^ of urease powder from *Canavalia ensiformis* (Jack bean). When fabricating silica nanomachines with attached peptides, different concentrations of AMP (500, 250, 125, 62.5, 31.25, 15.62, 7.81, 3.9, 1.95 and 0 μg mL^−1^) were added concomitantly with urease. The solution was kept mixing overnight (16 h) and then washed 3 times with 1x PBS (pH□=□7.4). After, the solution of urease nanomachines in 1x PBS (pH□=□7.4) was divided in aliquots and stored at 4 °C to be used for further experiments.

### Analysis of bioactive nanomachines motion

The Möbius from Wyatt Technology was used to analyze the electrophoretic mobility through dynamic light scattering (DLS) and extract both the apparent hydrodynamic radius and the diffusion coefficient of the urease nanomachines. Active motion of urease nanomachines was studied as a function of different concentrations of urea (500, 200, 50, 10, 1 and 0 mmol L^−1^) and AMPs (500, 250, 125, 62.5, 31.25, 15.62, 7.81, 3.9, 1.95 and 0 μg mL^−1^). The hydrodynamic radius is correlated with the diffusion coefficient according to the following Einstein-Stokes equation

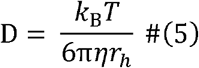

where *k*_*B*_ is the Boltzmann constant, *T* is the absolute temperature, *η* is the solvent viscosity and *r*_*h*_ is the hydrodynamic radius of the diffusing particle. The electrophoretic mobility was studied using an acquisition time of 5 s, with a laser of 532 nm wavelength and a detector angle of 163.5°. For each condition, the diffusion coefficient was calculated the average of 17-26 acquisitions obtained directly from the analysis of the scattering data on the Dynamics software. All the results are presented as mean ± standard error of the mean.

### Zeta potential analysis of bioactive micro- and nanomachines

The Zetasizer Nano S from Malvern Panalytical was used to study the electrophoretic mobility through dynamic light scattering (DLS) and obtain the zeta potential of the micro- and nanomachines. It was analyzed the zeta potential AMP-modified urease micromachines as a function of peptide concentration (500, 250, 125, 62.5, 31.25, 15.62, 7.81, 3.9 and 0 μg mL^−1^) and the zeta potential of AMP-modified urease nanomachines as a function peptide concentration (500, 250, 125, 62.5, 31.25, 15.62, 7.81, 3.9, 1.95 and 0 μg mL^−1^). For each condition, the result value was the mean of 3 measurements (which in its turn were an average of a minimum of 10 acquisitions), detected using a scattering angle of 173° and calculated using the Henry equation. The results are presented as mean ± standard error of the mean. The isoelectric points (pI) of the AMP were calculated through the web service Isoelectric Point Calculator (IPC) for peptides (http://isoelectric.org/).

### Bacterial strains and media

The strains used in this study were the pathogens *Escherichia coli* ATCC11775, *Acinetobacter baumannii* AB177, *Pseudomonas aeruginosa* PAO1, *Staphylococcus aureus* ATCC12600, *Klebsiella pneumoniae* ATCC13883. The bacteria were grown and plated on Luria-Bertani (LB) and Pseudomonas Isolation (*Pseudomonas aeruginosa* strains) agar plates incubated overnight at 37 °C. Following the incubation period, one isolated colony was transferred to 5 mL of medium (LB broth), which were incubated overnight (12–16 h) at 37 °C. In the following day, inoculums were prepared by diluting the bacterial overnight solutions 1:100 in 5 mL of the respective media and incubated at 37 °C until logarithmic phase (OD_600_ = 0.3-0.5).

### Antibacterial assays

Minimum inhibitory concentrations (MICs) of micro- and nanomachines were determined using the broth microdilution technique in LB with an initial inoculum of 5×10^6^ cells mL^−1^ in nontreated polystyrene microtiter plates (Corning, USA). Capsules were added to the plate as solutions in LB broth in concentrations ranging from 0 to 500 μg mL^−1^. The MIC was considered as the lowest concentration of the antimicrobial that completely inhibited the visible growth (readings were made in a spectrophotometer at 600 nm) of bacteria after 24 h of incubation of the plates at 37 °C. All assays were done in three independent replicates.

### Membrane depolarization assays

The cytoplasmic membrane depolarization activity of the AMP-urease machines was determined by measurements of fluorescence of the membrane potential–sensitive dye, DiSC_3_(5). Briefly, *A. baumannii* AB177 and *K. pneumoniae* ATCC13883 were grown at 37 °C with agitation until they reached mid-log phase (OD_600_= 0.5). The cells were then centrifuged and washed twice with washing buffer (20 mmol L^−1^ glucose, 5 mmol L^−1^ HEPES, pH 7.2) and re-suspended to an OD_600_ of 0.05 in the same buffer (20 mmol L^−1^ glucose, 5 mmol L^−1^ HEPES, pH 7.2), but containing 0.1 mol L^−1^ KCl. Thereafter, the cells (100 mL) were incubated for 15 min with 20 nmol L^−1^ of DiSC_3_(5) until a stable reduction of fluorescence was achieved, indicating the incorporation of the dye into the bacterial membrane. Membrane depolarization was then monitored by observing the change in the fluorescence emission intensity of the membrane potential–sensitive dye, DiSC_3_(5) (l_ex_ = 622 nm, l_em_ = 670 nm), after the addition of the micro- and nanomachines (100 mL solution at MIC values).

### Membrane permeabilization assay

The membrane permeability of the AMP-urease machines was determined by using the N-phenyl-1-napthylamine (NPN) uptake assay. *A. baumannii* AB177 and *K. pneumoniae* ATCC13883 were grown to an OD_600_ of 0.4, centrifuged (10,000 rpm at 4 °C for 10 min), washed and resuspended in buffer (5 mmol L^−1^ HEPES, 5 mmol L^−1^ glucose, pH 7.4). 4 mL of NPN solution (0.5 mmol L^−1^ – working concentration of 10 mmol L^−1^ after dilutions) was added to 100 mL of the bacterial solution in a white 96-wells plate. The background fluorescence was recorded at l_ex_= 350 nm and l_em_ = 420 nm. Micro- and nanomachines functionalized with urease and AMPs in water (100 μL solution at their MIC values) were added to the 96-wells plate, and fluorescence was recorded as a function of time until no further increase in fluorescence was observed (20 min).

### Skin abscess infection mouse model

*A. baumannii* AB177 was used to infect the murine skin. Briefly, bacteria were grown in tryptic soy broth medium. Subsequently, cells were washed twice with sterile PBS (pH 7.4, 13,000 rpm for 1 min), and resuspended to a final concentration of 5×10^6^ CFU/20 μL. Female CD-1 mice (six weeks-old) were anesthetized with isoflurane and had their backs shaved and a superficial linear skin abrasion was made with a needle to damage the stratum corneum and upper-layer of the epidermis. An aliquot of 20 μL containing the bacterial load in PBS was inoculated over each defined area containing the scratch with a pipette tip. Four hours after the infection, free peptides and micro and nanomachines functionalized with urease and AMPs at their MIC value were administered to the infected area. Animals were euthanized and the area of scarified skin was excised four-days post-infection, homogenized using a bead beater for 20 minutes (25 Hz), and 10-fold serially diluted for CFU quantification. Two independent experiments were performed with 8 mice per group in each condition.

## Acknowledgments

This work was supported the Spanish MINECO (project RTI2018-095622-B-I00), the Catalan AGAUR (project 2017 SGR 238), and the European Research Council (ERC) under the European Union’s Horizon 2020 research and innovation programme (grant agreement No 866348: iNanoSwarms). It was also funded by the grant RTI2018-098164-B-I00 from the MICIN/AEI/10.13039/5011000110333 and the “FEDER Una manera de hacer Europa” (BOTSinFluids project), the CERCA program by the Generalitat de Catalunya and the “Centro de Excelencia Severo Ochoa”, funded by Agencia Estatal de Investigación (CEX2018-000789-S). XA thanks the Spanish MINECO for the Severo Ochoa program (SEV-2014-0425) for the PhD fellowship (PRE2018-083712). CFN holds a Presidential Professorship at the University of Pennsylvania, is a recipient of the Langer Prize by the AIChE Foundation and acknowledges funding from the Institute for Diabetes, Obesity, and Metabolism, the Penn Mental Health AIDS Research Center of the University of Pennsylvania, the National Institute of General Medical Sciences of the National Institutes of Health under award number R35GM138201, and the Defense Threat Reduction Agency (DTRA; HDTRA11810041 and HDTRA1-21-1-0014). All figures were prepared using the Biorender drawing toolkit. The authors acknowledge the support of Ana Hortelão with the mesoporous silica particles synthesis and the TEM/SEM investigations.

## Authors’ contributions

SS and CFN conceived the idea and project. XA, TP and SS designed the micro- and nanomachines and conducted their fabrication, characterization, and motion experiments. XA performed the micro- and nanomachines fabrication, characterization, and motion experiments. MDTT and CFN designed all biological experiments. MDTT and AB performed all biological activity experiments, mechanism of action, and mouse work. XA, MDTT and CFN wrote the first draft of the manuscript. AB, TP, SS revised the manuscript. SS and CFN supervised and administered the work.

