## Supplementary Information for "Autonomous treatment of bacterial infections *in vivo* using antimicrobial micro- and nanomachines"

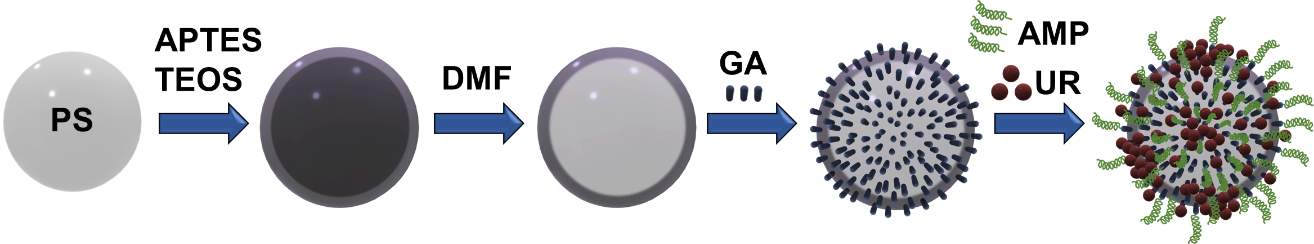

**Figure S1. Schematic of the synthesis of urease micromachines.** Polystyrene (PS) microbeads (D = 2 μm) were used to grow silica on top starting off 3-aminopropyltriethoxysilane (APTES) and tetraethylorthosilicate (TEOS) precursors. Dimethylformamide is used to dissolve the PS core and keep the silica shell where glutaraldehyde (GA) linker is used to attach urease (UR) and the antimicrobial peptide (AMP) payload.

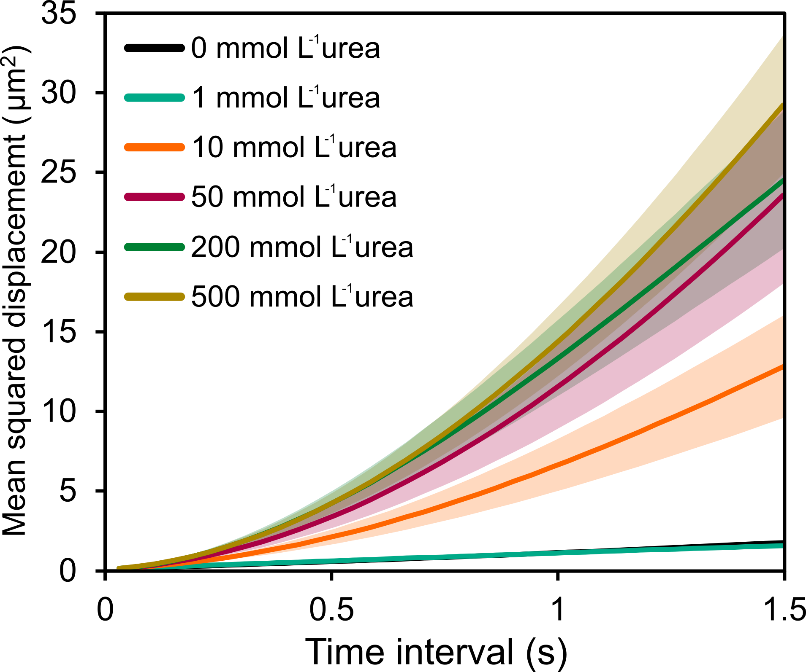

**Figure S2. Mean square displacement (MSD) of urea micromachines for urea concentration.** MSD growth extracted from X and Y position of micromachines recorded through microscopy, representing the average area explored for each time interval and under different concentrations of urea (substrate). All results are shown as the mean ± standard error of the mean.

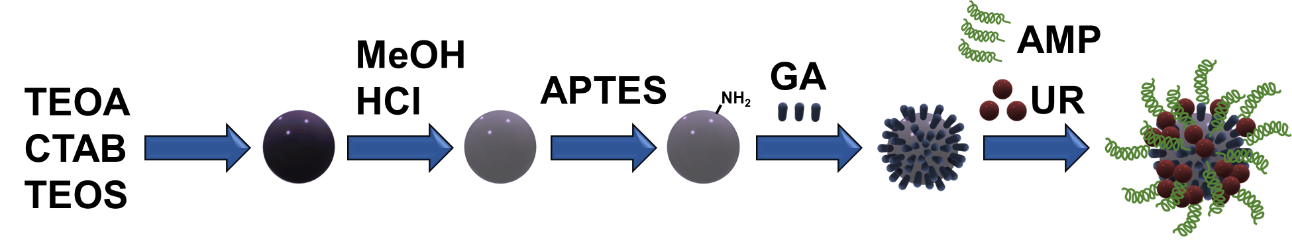

**Figure S3. Schematic of the synthesis of urease nanomachines.** Triethanolamine (TEOA) and hexadecyltrimethylammonium bromide (CTAB) are mixed to generate silica nanoparticles by dropwise addition of tetraethylorthosilicate (TEOS). After, the CTAB is removed by addition of methanol (MeOH) and hydrochloric acid (HCl) and the mesopores are formed. To functionalize the silica nanoparticles the surface is modified with amino groups adding 3-aminopropyltriethoxysilane (APTES). The glutaraldehyde (GA) linker is used to attach urease (UR) and the antimicrobial peptide (AMP) payload.

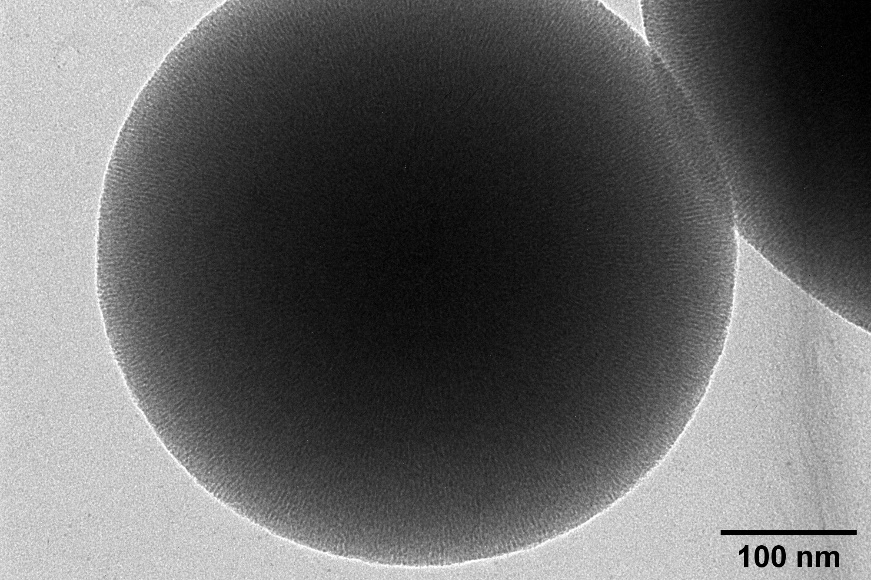

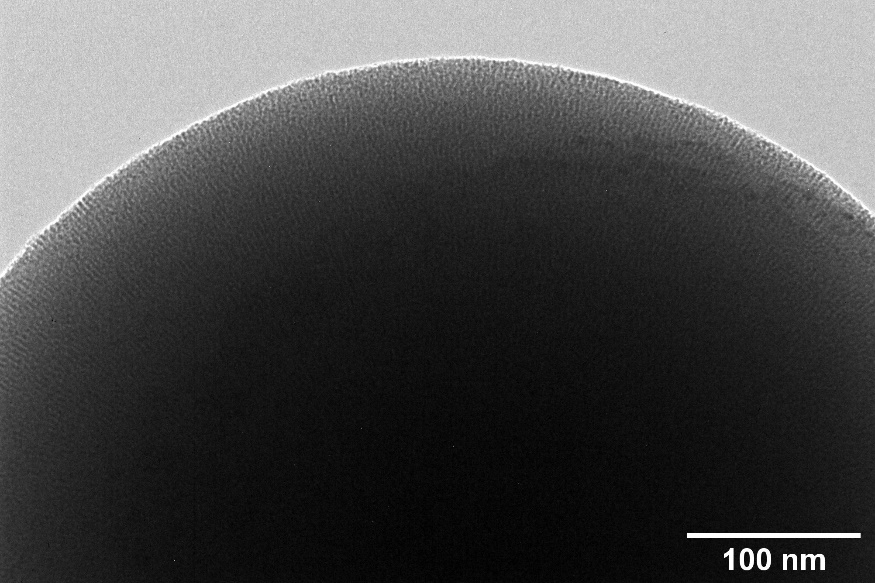

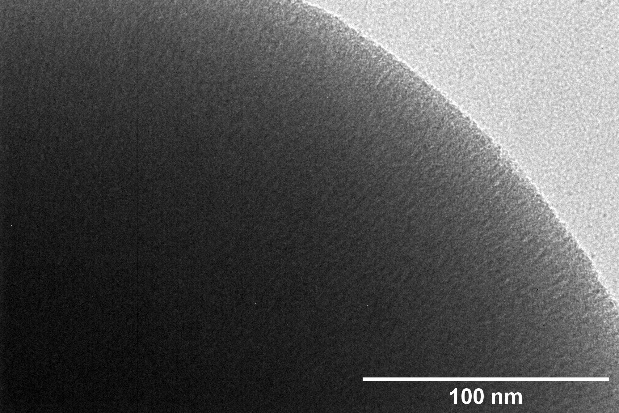

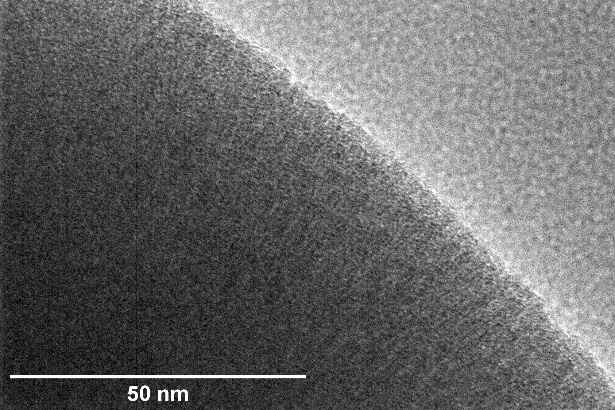

**Figure S4. TEM micrograph of silica nanoparticles (MSNP).** Micrographs of increasing magnification showing the radial mesoporosity channels inside the silica structure after the removal of CTAB, clearly visible specially in the outer area of the sphere.

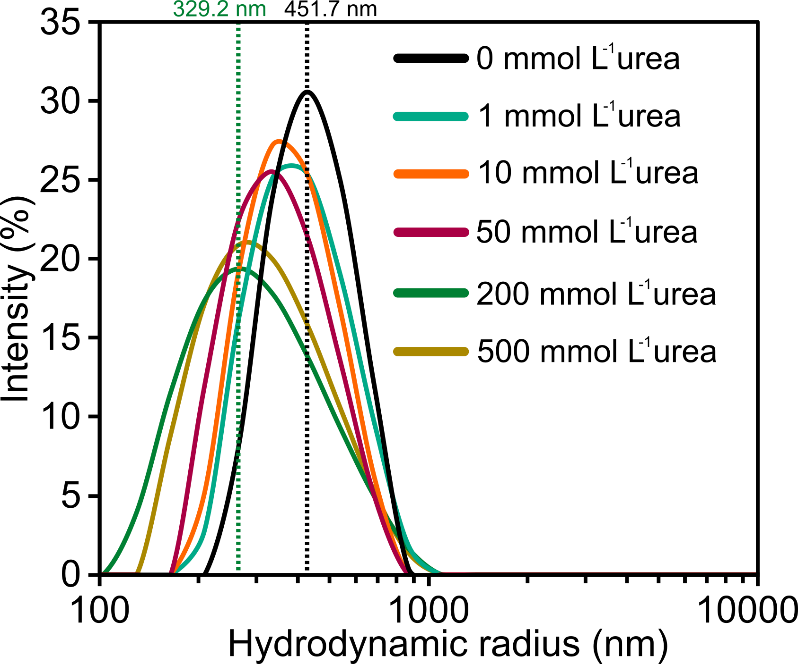

**Figure S5. Hydrodynamic radius of urease nanomachines for different concentrations of urea.** Representative hydrodynamic radii dispersion of urease nanomachines obtained through Dynamic Light Scattering (DLS) by measuring the diffusion of the particles and applying the Stokes−Einstein equation (see Methods section for details) to derive the radius for each urea concentration used.

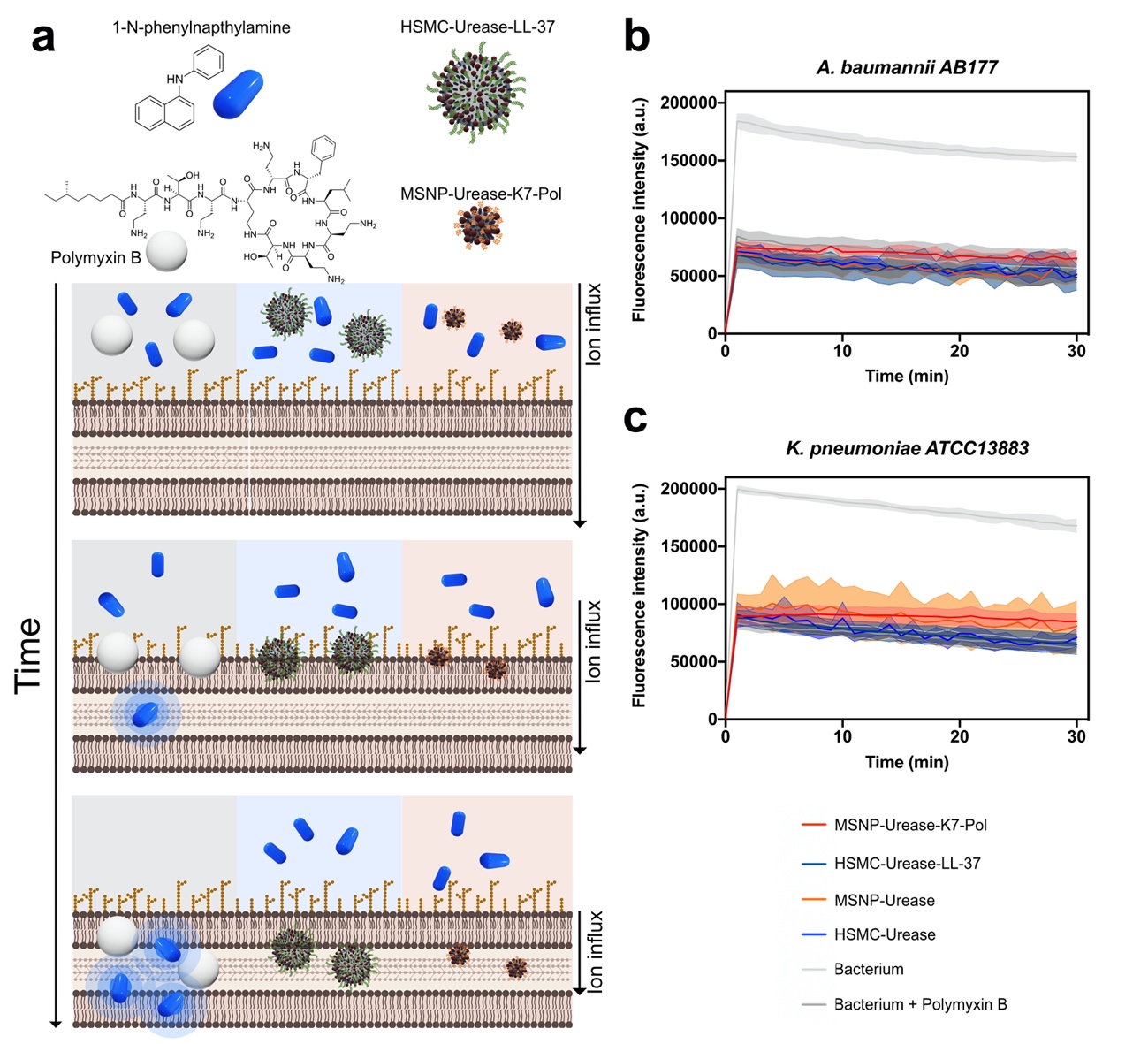

**Figure S6. Membrane permeabilization experiments with urease micro- and nanomotors functionalized with peptides.** **(a)** Motors do not permeabilize bacterial membranes at their MIC concentration against **(b)** *A. baumannii* AB177 and **(c)** *K. pneumoniae* ATCC13883. The permeabilizing antimicrobial polymyxin B was used as a positive control for permeabilization of the bacterial membranes.

**Figure S7. Images of the infection site on the back of mice used for the *in vivo* anti-infective model.** The images show (a) an untreated infection colonized by bacteria, (b) treatment with LL-37 that leads to partial wound healing, (c) treatment with K7-Pol that resolve the infection only at the administration site (bottom part of the wound), (d) treatment with the antimicrobial peptide payloads delivered by LL-37-urease micromachines that clear the whole extent of the wound and promote complete wound healing, and (e) treatment with the antimicrobial peptide payloads delivered by K7-Pol-urease nanomotors that resolve the infection but do not promote full wound healing.

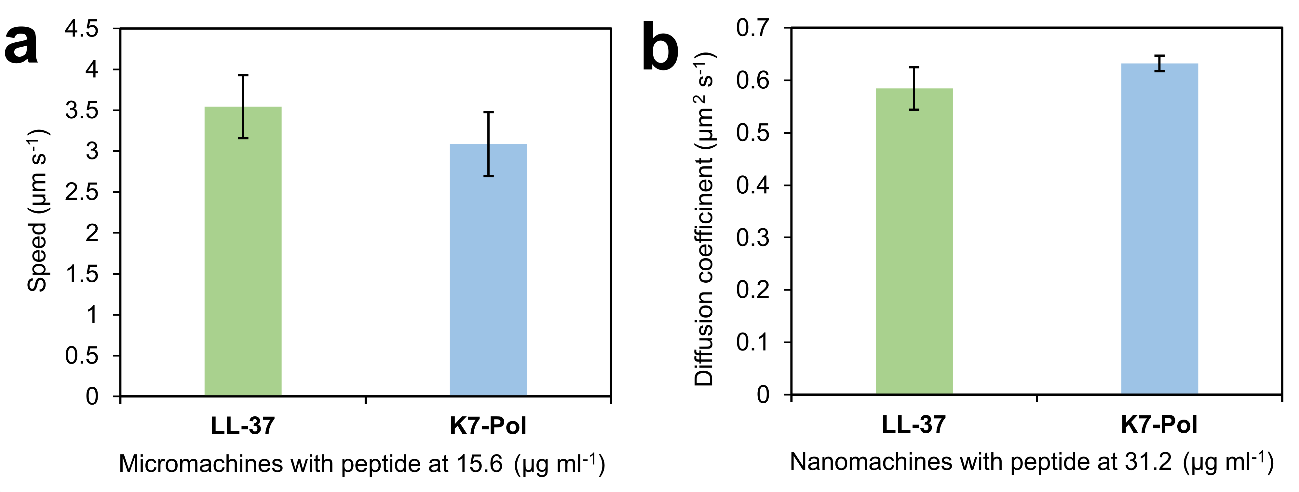

**Figure S8. Comparison of active motion of micro- and nanomachines with different antimicrobial peptides attached. (a)** Average speed of urease micromachines and **(b)** average diffusion coefficient of urease nanomachines modified with K7-Pol and LL-37 at the concentration used *in vivo* (15.6 μg mL^‒1^ for micromachines and 31.2 μg mL^‒1^ for nanomachines). No significant differences in motion were found between the peptides when anchored at the same concentration to the micro- and nanomachines.
